# Increased temperatures may safeguard the nutritional quality of crops under future elevated CO_2_ concentrations

**DOI:** 10.1101/344663

**Authors:** Iris H. Köhler, Steven C. Huber, Carl J. Bernacchi, Ivan R. Baxter

## Abstract

Iron (Fe) and zinc (Zn) deficiencies are a global human health problem that may worsen by growth of crops at elevated atmospheric CO_2_ concentration (eCO_2_). However, climate change will also involve higher temperature, but it is unclear how the combined effect of eCO_2_ and higher temperature will affect the nutritional quality of food crops. To begin to address this question, we grew soybean (*Glycine max*) in a Temperature by Free-Air CO_2_ Enrichment (T-FACE) experiment in 2014 and 2015 under ambient (400 *μ*mol mol^−1^) and elevated (600 *μ*mol mol^−1^) CO_2_ concentration and under ambient and elevated temperatures (+2.7 °C day and +3.4 °C at night). In our study, eCO_2_ significantly decreased Fe concentration in soybean seeds in both seasons (−8.7% and −7.7%) and Zn concentration in one season (−8.9%) while higher temperature (at ambient CO_2_ concentration) had the opposite effect. The combination of eCO_2_ with elevated temperature generally restored seed Fe and Zn concentrations to levels obtained under ambient CO_2_ and temperature conditions, suggesting that the potential threat to human nutrition by increasing CO_2_ concentration may not be realized. In general, seed Fe concentration was negatively correlated with yield suggesting inherent limitations to increasing seed Fe. In addition, we confirm our previous report that the concentration of seed storage products and several minerals varies with node position at which the seeds developed. Overall, these results demonstrate the complexity of predicting climate change effects on food security when various environmental parameters change in an interactive manner.

## Significance Statement

Mineral deficiencies are a global health problem that may be exacerbated by crop growth at elevated atmospheric CO_2_ concentration (eCO_2_) that can reduce micronutrient concentrations in grains and seeds. However, our soybean studies suggest that the potential threat to human nutrition by increasing CO_2_ concentration may not be realized when plants grow at eCO_2_ and higher temperature that more realistically mimic future climate conditions.

## Introduction

Atmospheric CO_2_ is increasing and projected to reach between 730 and 1020 μmol mol^−1^ by the end of the century (Collins et al., 2013). The increase in CO_2_ would be expected to increase leaf photosynthesis and this has been documented experimentally for many C_3_ species (Ainsworth and Long, 2005, Ainsworth and Rogers, 2007) using free-air CO_2_ enrichment (FACE) technology (Long et al., 2004, Leakey et al., 2009). As a result, biomass production and seed yield are increased but often to a lesser extent than the increase in light-saturated photosynthesis (Long et al., 2004, Leakey et al., 2009). In addition to increasing yield, eCO_2_ has the potential to reduce the concentration of minerals in seeds and thereby threaten human nutrition, as highlighted in two recent meta-analysis studies. Across a range of C_3_ plants, Loladze (Loladze, 2014) reported that eCO_2_ reduced the concentrations of several minerals in foliar and edible tissues by approximately 8%, with the exception of Mn that was not reduced. Overall, the patterns of mineral changes were similar between foliar and edible tissue with the exception that K was reduced only in edible tissues (Loladze, 2014). The second study (Myers et al., 2014) focused on C_3_ grains and legumes and reported that concentrations of Zn and Fe were decreased from ~5 to 10% at eCO_2_. Both studies concluded that eCO_2_ may result in crops that are less nutritious as a result of reduced mineral concentrations, which would have enormous implications for the use of crops for human food as well as animal feed (Weigel, 2014).

The mechanisms responsible for the effects of eCO_2_ on seed mineral concentrations are not clear, but many biological and physical processes could contribute. For example, reduced transpiration as a result of partial stomatal closure at eCO_2_ (Bernacchi et al., 2007, Zaman et al., 2013) would be expected to reduce the uptake of minerals that are acquired by mass flow, but have less effect on those where movement to the root surface occurs by diffusion (McGrath and Lobell, 2013). Following uptake into the plant, mineral transport continues in the xylem with distribution among stems and leaves (Marschner, 1995). In grasses, nutrient distribution among vegetative organs is known to be controlled independently of transpiration rate (Yamaji and Ma, 2017), and it is likely that this occurs in dicots as well. The mineral content of seeds is thought to reflect import from the phloem as a result of continued uptake by roots (followed by xylem-to-phloem transfer of minerals) as well as redistribution from vegetative tissues such as leaves. Conceivably, many of these steps could be impacted by eCO_2_. Furthermore, dilution by enhanced carbohydrate production at eCO_2_ or dilution by increased seed production could contribute to the reduced mineral concentrations observed. Dilution by enhanced carbohydrate production cannot explain all of the effect of eCO_2_ (McGrath and Lobell, 2013), but the impact of changes in seed yield on the content of minerals on a plant basis (Singh et al., 2016) have not been broadly considered. However, it is reasonable to assume that if a micronutrient is obtained in controlled or limited amounts it would be diluted when distributed amongst a greater number of seeds produced at eCO_2_, such that the concentration would decrease while the mineral seed content per plant would not change.

Another environmental factor that needs to be considered along with eCO_2_ is elevated temperature. It is generally recognized that an increase in atmospheric CO_2_, coupled with atmospheric accumulation of other greenhouse gases, will be accompanied by an increase in terrestrial surface air temperatures between 1 to 6 °C by 2050, relative to 1961–1990, depending on geographic location (Rowlands et al., 2012). The impact of elevated temperature would be expected to have a somewhat greater effect on reproductive development, as soybean has an optimum temperature for vegetative growth of ~30 °C (Hesketh et al., 1973) whereas reproductive development has an optimal temperature of 22 to 24 °C (Hatfield et al., 2011). Furthermore, an increase in temperature will generally lead to an increase in vapor pressure deficit (VPD) that would increase transpiration (assuming water supply is sufficient) and as a result the uptake of minerals driven by mass flow. Thus, elevated temperature could impact the concentration of minerals in plant tissues including seeds and in an opposite manner to eCO_2_.

Because future climate change is projected to involve both rising CO_2_ and temperature it is important to evaluate both environmental factors together. Recently, the Temperature by Free-Air CO_2_ Enrichment (T-FACE) facility was used to explore the independent and interactive effects of eCO_2_ and elevated temperature on soybean photosynthesis and productivity (Ruiz-Vera et al., 2013, Köhler et al., 2016). In general, photosynthesis, aboveground biomass, and seed yields were increased by eCO_2_ and decreased by elevated temperature; the combination of eCO_2_ plus elevated temperature had a somewhat variable effect (likely dependent on ambient temperature during the growing season) but the generalized conclusion was that eCO_2_ attenuated the negative impact of elevated temperature (Ruiz-Vera et al., 2013). However, in terms of seed mineral concentrations, there is little information about the impact of eCO_2_ in combination with elevated temperature. Consequently, the overall objective of the present study was to determine the individual and combined effect of eCO_2_ and elevated temperature on soybean seed composition in terms of storage products (protein and oil) and important minerals. Furthermore, we examined seed produced at different positions along the main stem because seed composition varies with nodal position (Huber et al., 2016). Overall, the results suggest that eCO_2_ in combination with elevated temperature may reduce seed yield relative to eCO_2_ conditions but will safeguard the mineral nutrition quality of soybeans.

## Results

### Plant Growth and Seed Yield

Soybean plants were grown in the field under conditions of eCO_2_ and elevated temperature similar to predicted conditions in 2050. In both seasons of our study (2014 and 2015), eCO_2_ increased plant height (Figure 1a) and the number of main stem nodes (Figure 1b) compared to plants at atmospheric [CO_2_], whereas elevated temperature had the opposite effect but only in 2014 (Table 1).

In both years eCO_2_ significantly increased total seed yield while elevated temperature significantly decreased total seed yield in the bulk samples collected from the 3.2 m row harvest (Figure 2, Table 1). A similar pattern was observed for the total seed yield from the replicated subsampling of 10 single plants, and yield data from both sampling methods were closely correlated (R^2^ = 0.82). The 10-plant samples consisted of normal sized, representative plants selected for each of the treatments that were harvested at maturity and the main stems divided into thirds, with each stem fraction (bottom, middle, and top) threshed separately. The distribution of seed yield among the three main stem positions is presented in Figure 3a and as a percentage distribution in Figure 3b. As expected, 40 to 50% of the total main stem seed yield was located in the middle canopy position in both years. In 2014 a higher proportion (25% to 33%) of the total seed yield was located at the bottom canopy position compared to 2015 (13% to 23%), and consequently the opposite was observed for the proportion at the top canopy position in 2014 (24% to 33%) and 2015 (30% to 42%) (Figure 3b). Elevated CO_2_ affected the proportional distribution of seed yield between the three canopy positions, but only in 2015, whereas the heat treatment led to a decreased seed yield at the bottom canopy position and increased seed yield at the top canopy position under both ambient and eCO_2_ concentrations in both seasons (Figure 3b, Table 2).

**Figure 1.**
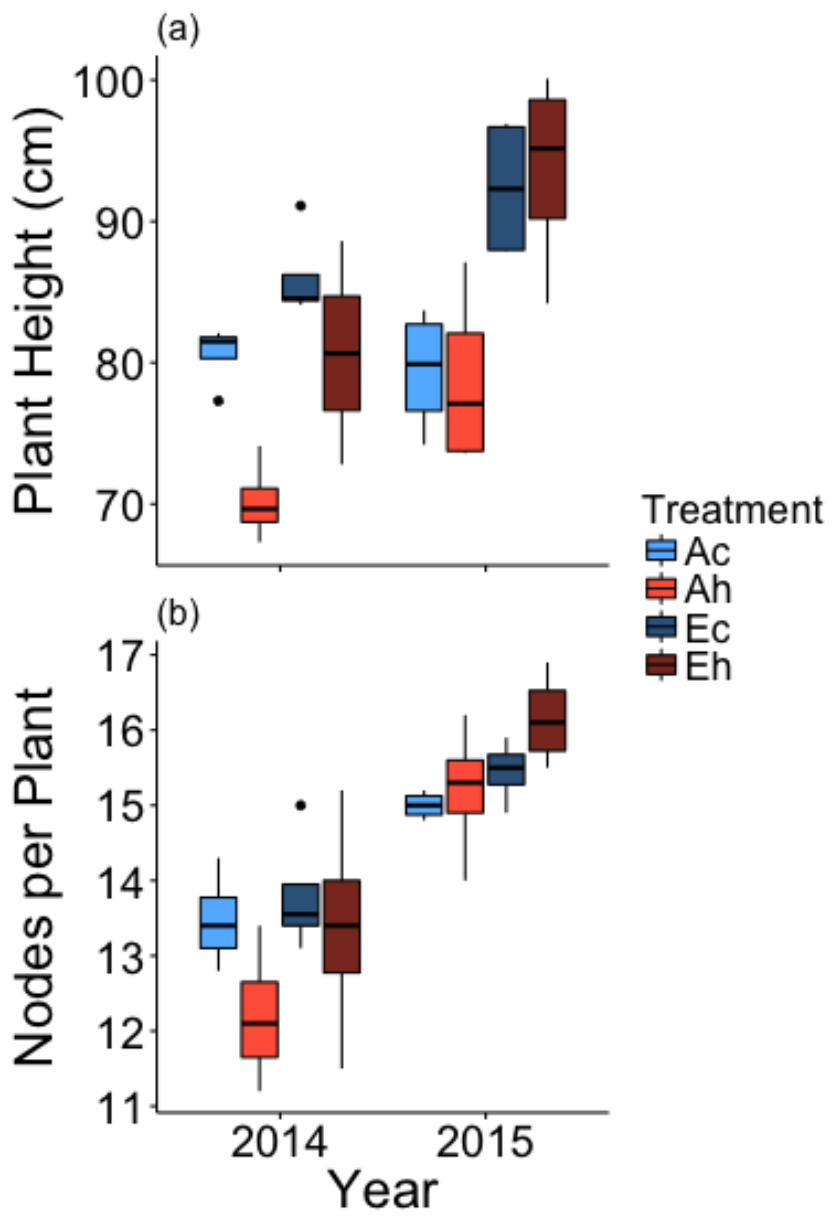
Impact of eCO_2_ and elevated temperature treatments, singly and in combination, on (a) mean plant height. and (b) number of main stem nodes per plant for soybean over two growing seasons. Both the CO_2_ (F1,9 = 15.95; p< 0.005) and temperature (F1,9 = 15.66; p < 0.005) main effects were statistically significant in 2014. The CO_2_ main effect (F1,9 = 28.23) was statistically significant in 2015. There were no statistically significant interactions between CO_2_ and temperature for either height or node number in either year. Boxplots display the median, and 25 and 75% percentiles, and whiskers extend to 1.5X interquartile range, with outliers (if any) shown.

**Figure 2:**
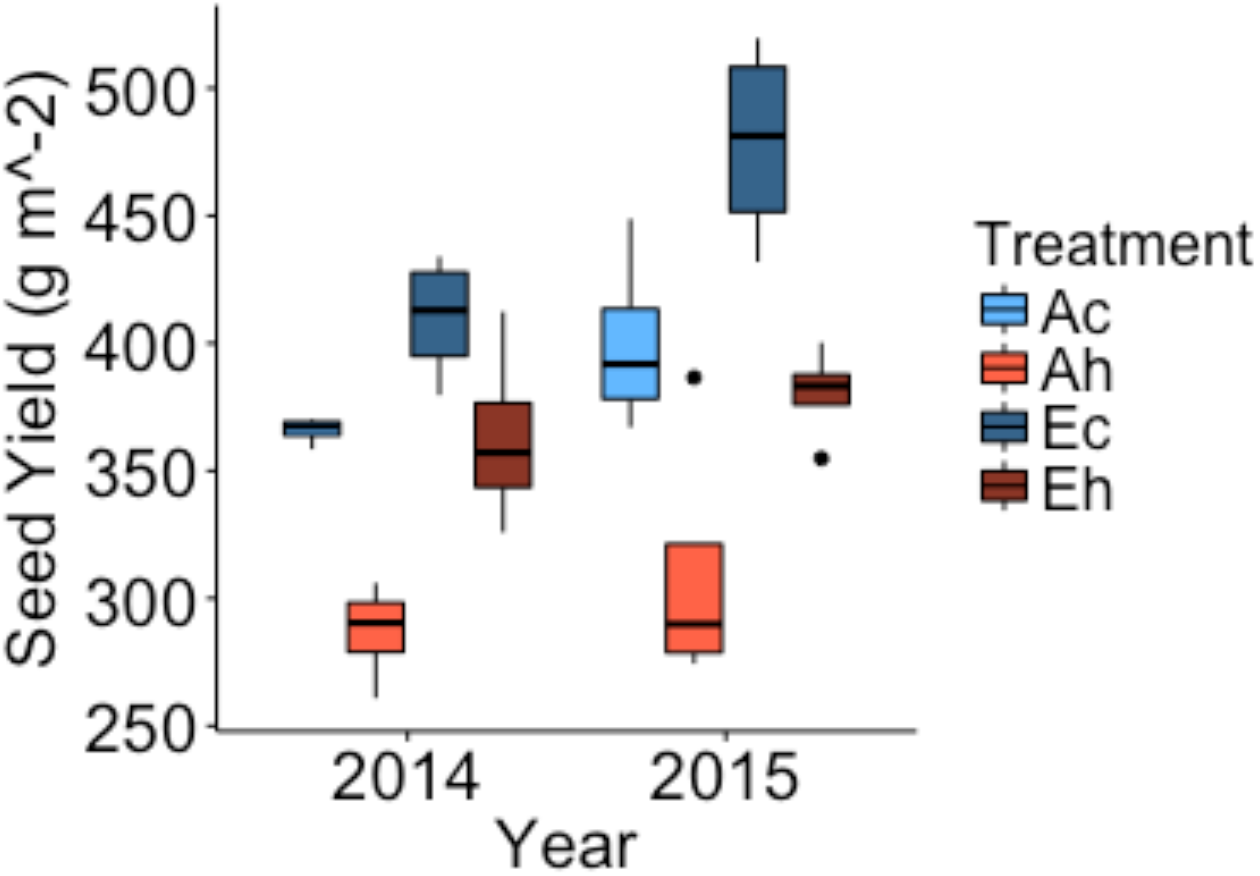
Impact of eCO_2_ and elevated temperature treatments, singly and in combination, on seed yield in g m^−2^ as derived from sampling of 3.2m rows. Ac, ambient CO_2_, control temperature; Ah, ambient CO_2_, heated + 3.5 °C; Ec, elevated CO_2_, control temperature; Eh, elevated CO_2_, heated + 3.5 °C) in 2014 and 2015. Boxplots display the median, and 25 and 75% percentiles, and whiskers extend to 1.5X interquartile range, with outliers (if any) shown.

**Table 1:** Complete block analysis of variance of total seed yield (SY) for the main effects [CO_2_] concentration (400 μmol mol^−1^, 600 μmol mol^−1^), temperature (control, heat) and the interaction term. Values in table are *p*-values, significance was set as *p* < 0.1.

|  | Seed Yield |  |  |
| --- | --- | --- | --- |
|  | CO<sub>2</sub> | Temp | CO<sub>2</sub>*<br>Temp |
| 2014 | **0.0231** | **0.0003** | 0.1055 |
| 2015 | **0.0192** | **<.0001** | 0.688 |
| *Num DF* | 1 | 1 | 1 |
| *Den DF* | 3 | 6 | 6 |
|  | Plant Height |  |  |
|  | CO<sub>2</sub> | Temp | CO<sub>2</sub>*<br>Temp |
| 2014 | **0.031** | **0.0033** | 0.2394 |
| 2015 | **<0.001** | **0.9176** | 0.7104 |
| *Num DF* | 1 | 1 | 1 |
| *Den DF* | 9 | 9 | 9 |
|  | Nodes per Plant |  |  |
|  | CO<sub>2</sub> | Temp | CO<sub>2</sub>*<br>Temp |
| 2014 | **0.0574** | **0.0311** | 0.2568 |
| 2015 | **0.0124** | **0.0746** | 0.2689 |
| *Num DF* | 1 | 1 | 1 |
| *Den DF* | 9 | 9 | 9 |

Although there was some variation in single seed weight from the samples collected at different canopy positions in the present study (Figure S1 and Köhler et al., 2016), these differences were generally small. Across all treatments, comparison of single seed weight with seed yield at the corresponding canopy position (i.e., bottom, middle or top position) produced a positive correlation but with a very low correlation coefficient (r = 0.28) that was not statistically significant. When different environmental parameters (i.e., elevated temperature or eCO_2_) or canopy positions were color-coded, it was apparent that the weak correlation was valid across all of the parameters (Figure S2). The implication is that number of seeds produced rather than changes in seed size primarily drove changes in yield.

### Seed Protein and Oil Concentrations

Canopy position significantly affected seed composition of major storage products, with higher oil concentrations (Figure 4a,b) at the bottom of the canopy and conversely, higher protein concentrations (Figure 4c,d) at the top of the canopy. In general, the canopy gradients in protein and oil concentration were larger in 2015 compared to 2014. There was no significant effect of eCO_2_ on protein (Figure 4c,d) or oil (Figure 4a,b) concentrations in the three canopy layers (Table 3). In contrast, elevated temperature significantly reduced protein concentration in 2014 (Figure 4c) with the strongest effect (−1.6%, p < 0.0001) in both the bottom and middle canopy positions and a smaller effect at the top of the canopy (−0.6%, p < 0.01) whereas in 2015, a significant reduction was only observed at the middle canopy position (−0.9%, p < 0.1). Conversely, the elevated temperature treatment significantly increased oil concentration at all positions in 2014 (bottom: +1.0%, p < 0.0001; middle: +0.9%, p < 0.0001, and top: +0.8%, p < 0. 0001) and at the bottom (+0.5%, p < 0.05) and middle position (+0.5%, p < 0.05) in 2015.

**Figure 3:**
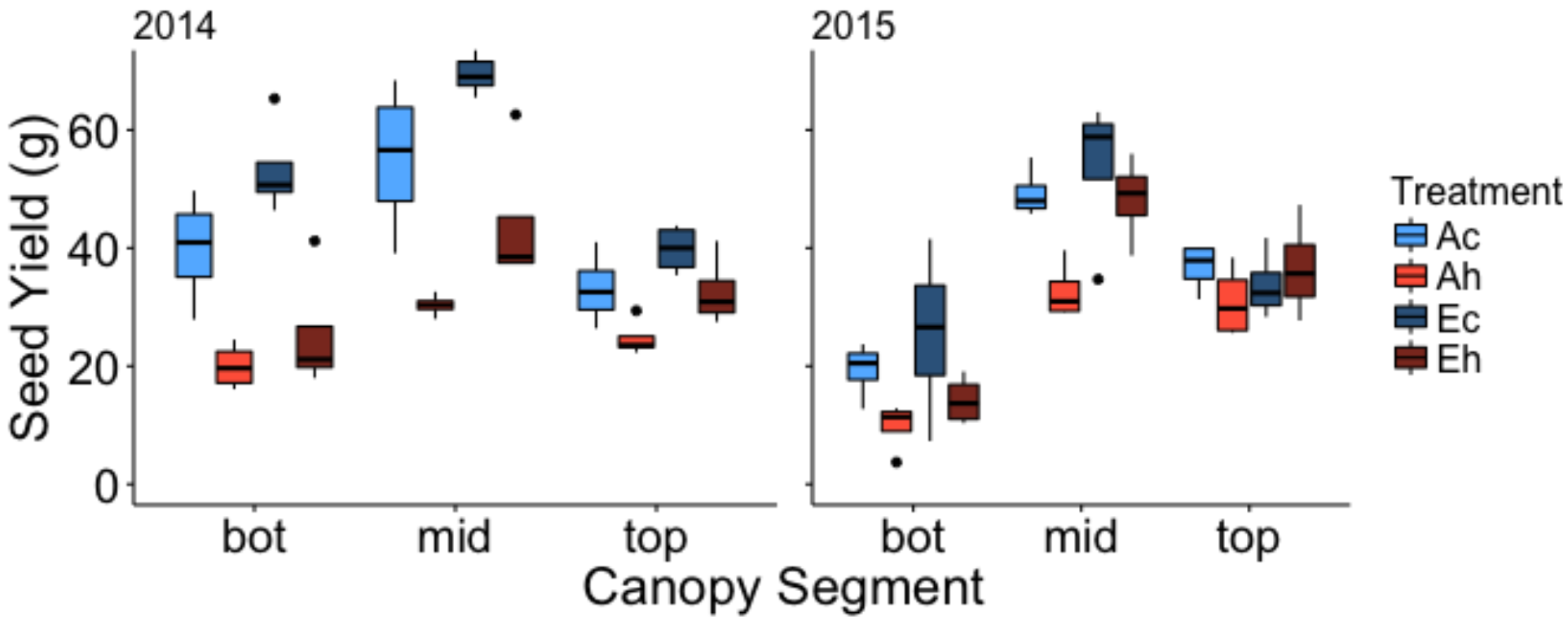
Distribution of seed yield among the bottom, middle and top canopy positions as impacted by eCO_2_ and elevated temperature treatments in 2014 and 2015. (a) Absolute seed yield expressed as g from each 10-plant sample. (b) Seed yield from each position expressed as a percentage of total seed under the four treatments. Ac, ambient CO_2_, control temperature; Ah, ambient CO_2_, heated + 3.5 °C; Ec, elevated CO_2_, control temperature; Eh, elevated CO_2_, heated + 3.5 °C. Boxplots display the median, and 25 and 75% percentiles, and whiskers extend to 1.5X interquartile range, with outliers (if any) shown.

**Table 2:** Complete block analysis of variance of the SY (g) data from the 10 plants samples for the main effects [CO_2_] concentration (400 μmol mol^−1^, 600 μmol mol^−1^), temperature (control, heat), canopy position (bottom, middle, top) and the interaction terms. Values in table are *p*-values, significance was set as *p* < 0.1.

|  | CO<sub>2</sub> | Temp | CO<sub>2</sub>*<br>Temp | Canopy | CO<sub>2</sub>*<br>Canopy | Temp*<br>Canopy | CO<sub>2</sub>*<br>Temp*<br>Canopy |
| --- | --- | --- | --- | --- | --- | --- | --- |
| Seed Yield (g) |  |  |  |  |  |  |  |
| 2014 | **0.035** | 0.2657 | 0.7855 | **<0.0001** | **0.0966** | **0.0003** | 0.3225 |
| 2015 | 0.2323 | **0.0043** | 0.1293 | **<0.0001** | 0.1349 | **0.058** | 0.2827 |
| Seed Yield (%) |  |  |  |  |  |  |  |
| 2014 | - | - | - | **<0.0001** | 0.4593 | **<0.0001** | 0.3329 |
| 2015 | - | - | - | **<0.0001** | **0.0988** | **0.0065** | 0.7987 |
| *Num DF* | 1 | 1 | 1 | 2 | 2 | 2 | 2 |
| *Den DF* | 3 | 6 | 6 | 24 | 24 | 24 | 24 |

**Figure 4:**
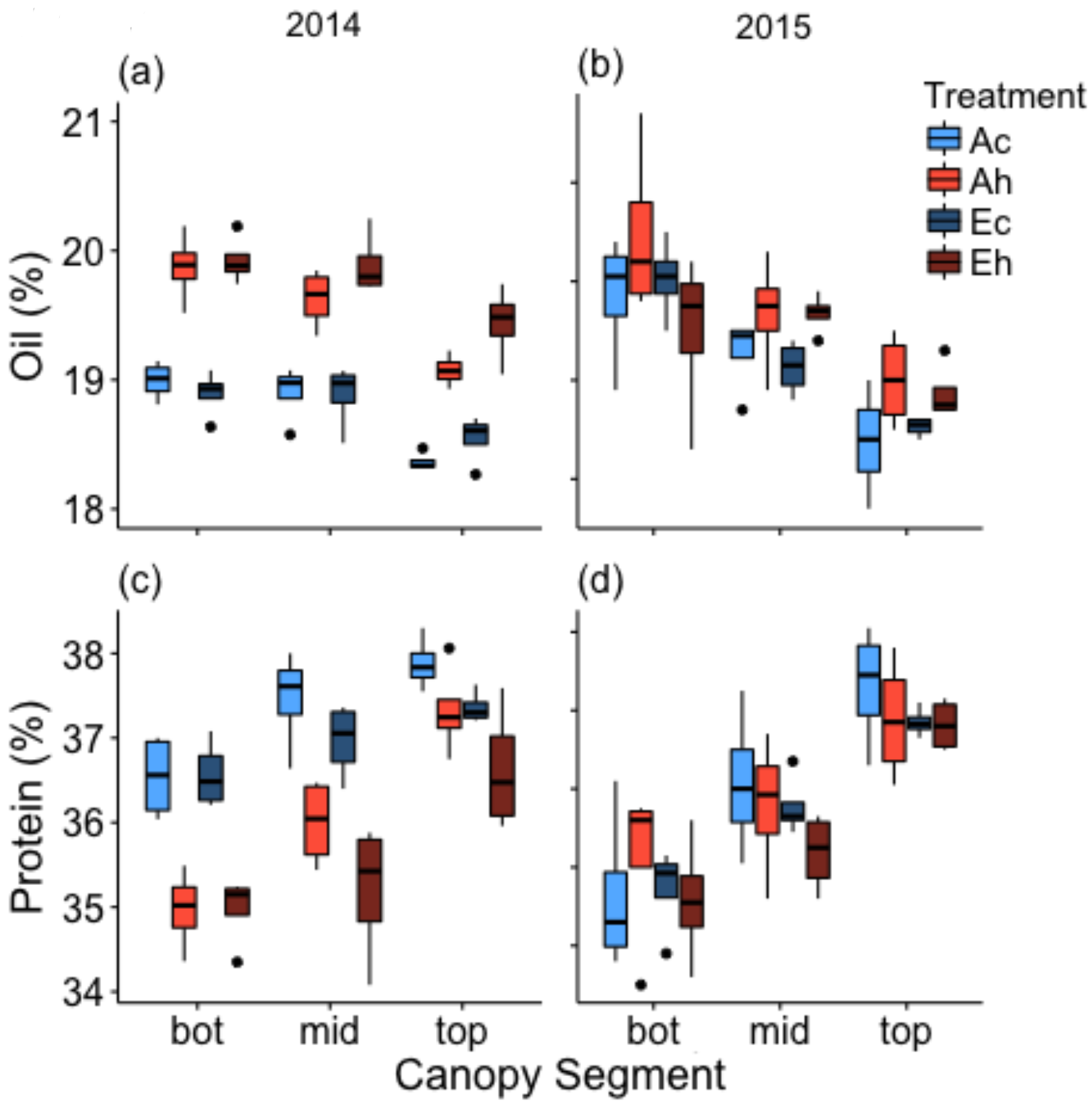
Concentration of (a,b) oil and (c,d) protein in seeds produced at the bottom, middle and top canopy positions as impacted by eCO_2_ and elevated temperature treatments in 2014 and Ac, ambient CO_2_, control temperature; Ah, ambient CO_2_, heated + 3.5 °C; Ec, elevated CO_2_, control temperature; Eh, elevated CO_2_, heated + 3.5 °C. Boxplots display the median, and 25 and 75% percentiles, and whiskers extend to 1.5X interquartile range, with outliers (if any) shown.

**Table 3:** Complete block analysis of variance of the seed protein and oil concentrations in the two seasons for the main effects [CO_2_] concentration (400 μmol mol^−1^, 600 μmol mol^−1^), temperature (control, heat), canopy position (bottom, middle, top) and the interaction terms. Values in table are *p*-values, significance was set as *p* < 0.1.

|  | CO<sub>2</sub> | Temp | CO<sub>2</sub>*<br>Temp | Canopy | CO<sub>2</sub>*<br>Canopy | Temp*<br>Canopy | CO<sub>2</sub>*<br>Temp*<br>Canopy |
| --- | --- | --- | --- | --- | --- | --- | --- |
| Protein |  |  |  |  |  |  |  |
| 2014 | 0.2192 | <.0001 | 0.5433 | <.0001 | **0.0381** | **0.0018** | 0.8788 |
| 2015 | 0.5339 | 0.3689 | 0.6311 | <.0001 | 0.6459 | **0.0646** | 0.1477 |
| Oil |  |  |  |  |  |  |  |
| 2014 | 0.2338 | <.0001 | 0.2531 | <.0001 | **0.0524** | 0.419 | 0.8494 |
| 2015 | 0.6046 | **0.0829** | 0.2419 | <.0001 | 0.3068 | 0.1657 | **0.078** |
| *Num DF* | 1 | 1 | 1 | 2 | 2 | 2 | 2 |
| *Den DF* | 3 | 6 | 6 | 24 | 24 | 24 | 24 |

### Seed mineral composition

Seeds of legumes such as soybeans are generally considered a good source of micronutrients (minerals) that are important for human health. Consequently, it is important that environmental factors that affect the concentrations of minerals in seeds are identified, and as noted above, eCO_2_ has been reported to reduce the concentrations of important minerals in soybean seeds and other edible grains (Loladze, 2014, McGrath and Lobell, 2013, Myers et al., 2014). A significant effect of eCO_2_ on mineral concentrations was observed in the present study for six elements (B, Ca, Fe, K, Se, Zn), but was only consistently observed in both study years for Fe. In contrast, elevated temperature had a significant effect on 14 of the analyzed elements in at least one year of the study and this effect was observed in both study years for 7 of those elements (As, B, Ca, Fe, Ni, Rb, Sr). In some cases, significant interactions were also observed, in particular between temperature and canopy position. Consistent with our earlier report (Huber et al., 2016), canopy position had a significant effect on seed elemental concentrations of 17 of the 20 analyzed elements in at least one year of the study and for 10 of those elements (Ca, Co, Cu, Fe, Mg, Mn, P, Rb, Sr, Zn) the effect was observed in both years of the study. For a summary of the results, see Tables 4 and 5. In the following, we present the results for nutritionally important minerals [e.g., Fe, Zn, Mg, Ca, Se and Cu; (USDA, 2015)] and others of particular interest in more detail below.

#### Iron and magnesium

Both minerals are important for human nutrition and varied with canopy position in the same way, with higher concentrations in seeds produced at the bottom compared to the top of the canopy (Figure 5). However, only the seed concentration of iron [Fe] was affected by eCO_2_ and elevated temperature in both seasons. Seed [Fe] was significantly lower under eCO_2_ at all canopy positions (Figure 5a). On average, eCO_2_ decreased seed [Fe] by 8.7% in 2014 and by 7.7% in 2015. In contrast, elevated temperature significantly increased [Fe] by 11.1% in 2014 and by 7.4% in 2015 over the control temperature treatment irrespective of [CO_2_]. In both years and under all treatments, seed [Fe] decreased towards the top of the canopy but in 2015 this effect was more pronounced in the control temperature treatments, with −23.2% lower [Fe] at the top compared to the bottom, while under the heat treatment this decrease was only −13%.

**Table 4:** Complete block analysis of variance of the elemental mineral concentrations in soybean seeds in the two seasons for the main effects [CO_2_] concentration (400 μmol mol^−1^, 600 μmol mol^−1^), temperature (control, heat), canopy position (bottom, middle, top) and the interaction terms.

| Element | Year | CO<sub>2</sub> | Temp | CO<sub>2</sub>*<br>Temp | Canopy | CO<sub>2</sub>*<br>Canopy | Temp*<br>Canopy | CO<sub>2</sub>*<br>Temp*<br>Canopy |
| --- | --- | --- | --- | --- | --- | --- | --- | --- |
| *Al* | 2014 | ns | ns | ns | ns | ns | ns | ns |
|  | 2015 | ns | <0.05 | ns | ns | ns | ns | ns |
| *As* | 2014 | ns | <0.05 | ns | ns | ns | ns | ns |
|  | 2015 | ns | <0.1 | ns | ns | ns | ns | ns |
| *B* | 2014 | ns | <0.001 | ns | <0.1 | ns | <0.0001 | ns |
|  | 2015 | <0.1 | <0.0001 | <0.1 | ns | <0.01 | <0.0001 | <0.05 |
| *Ca* | 2014 | <0.1 | <0.01 | ns | <0.0001 | ns | ns | ns |
|  | 2015 | ns | <0.01 | ns | <0.01 | ns | <0.05 | ns |
| *Cd* | 2014 | ns | <0.1 | ns | ns | ns | <0.01 | ns |
|  | 2015 | ns | ns | ns | ns | ns | <0.1 | ns |
| *Co* | 2014 | ns | <0.05 | <0.1 | <0.0001 | ns | <0.001 | ns |
|  | 2015 | ns | ns | ns | <0.001 | ns | <0.05 | ns |
| *Cu* | 2014 | ns | ns | ns | <0.0001 | <0.01 | <0.01 | <0.1 |
|  | 2015 | ns | ns | ns | <0.0001 | <0.05 | <0.01 | ns |
| *Fe* | 2014 | <0.1 | <0.001 | ns | <0.0001 | ns | ns | ns |
|  | 2015 | <0.05 | <0.01 | ns | <0.0001 | ns | <0.05 | ns |
| *K* | 2014 | <0.1 | ns | ns | ns | <0.05 | ns | ns |
|  | 2015 | ns | ns | ns | <0.0001 | ns | ns | <0.1 |
| *Mg* | 2014 | ns | ns | ns | <0.01 | ns | ns | ns |
|  | 2015 | ns | <0.1 | ns | <0.05 | ns | <0.1 | ns |
| *Mn* | 2014 | ns | ns | ns | <0.0001 | ns | ns | <0.1 |
|  | 2015 | ns | <0.1 | ns | <0.0001 | ns | ns | ns |
| *Mo* | 2014 | ns | ns | ns | <0.0001 | <0.05 | ns | ns |
|  | 2015 | ns | ns | ns | ns | ns | ns | ns |
| *Na* | 2014 | ns | ns | ns | <0.01 | ns | ns | ns |
|  | 2015 | ns | ns | ns | ns | ns | ns | ns |
| *Ni* | 2014 | ns | <0.05 | ns | ns | ns | <0.01 | ns |
|  | 2015 | ns | <0.01 | ns | <0.05 | ns | ns | ns |
| *P* | 2014 | ns | <0.001 | ns | <0.05 | <0.05 | <0.05 | ns |
|  | 2015 | ns | ns | ns | <0.1 | ns | <0.1 | ns |
| *Rb* | 2014 | ns | <0.01 | ns | <0.05 | ns | <0.01 | ns |
|  | 2015 | ns | <0.05 | ns | <0.0001 | ns | ns | ns |
| *S* | 2014 | ns | <0.1 | ns | ns | ns | <0.05 | ns |
|  | 2015 | <0.05 | ns | ns | <0.0001 | ns | ns | ns |
| *Se* | 2014 | ns | ns | <0.1 | <0.1 | <0.05 | ns | ns |
|  | 2015 | ns | ns | <0.1 | ns | ns | <0.05 | ns |
| *Sr* | 2014 | ns | <0.05 | ns | <0.0001 | <0.05 | ns | ns |
|  | 2015 | ns | <0.01 | ns | <0.0001 | ns | <0.1 | ns |
| *Zn* | 2014 | ns | ns | ns | <0.1 | ns | <0.05 | ns |
|  | 2015 | <0.1 | ns | ns | <0.01 | ns | <0.1 | ns |
| *Num DF* |  | 1 | 1 | 1 | 2 | 2 | 2 | 2 |
| *Den DF* |  | 3 | 6 | 6 | 24 | 24 | 24 | 24 |

**Table 5:**
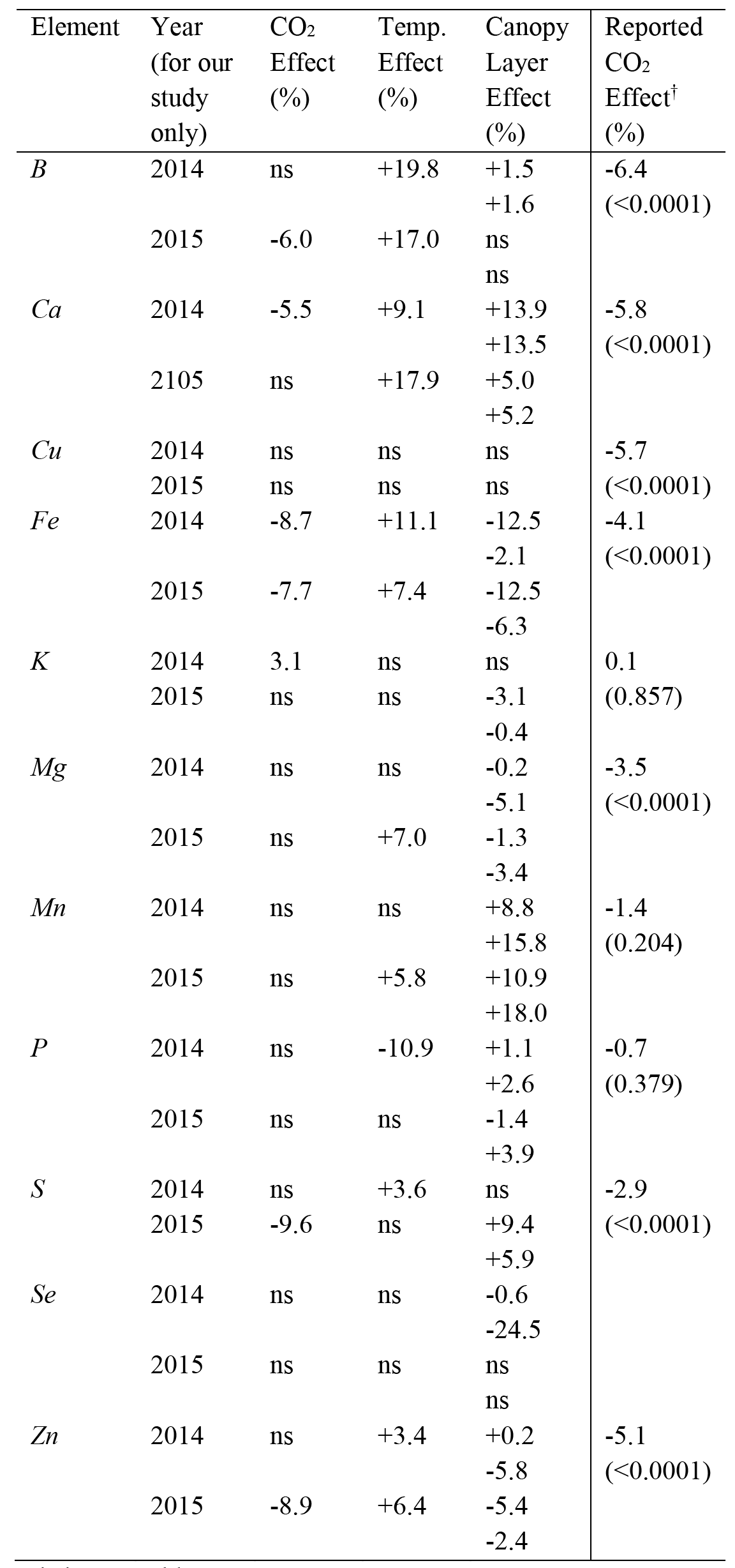
Overview of CO_2_, temperature and canopy layer effects (magnitude and direction) for selected mineral elements measured in this study and reported CO_2_ effects. Values reported for the canopy layer effect are middle *vs*. bottom layer (first row) and top *vs*. middle layer (second row). For the respective p-values see Table 4.

Magnesium concentration [Mg] was not significantly affected by eCO_2_ (Figure 5b). Seed [Mg] decreased towards the top of the canopy in 2014 (top *vs* bottom: −5.2%) and 2015 (top *vs* bottom: −4.7%). In 2015, higher seed [Mg] was observed under the heat treatment (+7.0%) in comparison to the control temperature treatment and this effect was larger in seeds at the top of the canopy (+11.9%) compared to the middle (+4.2%) or bottom canopy (+5.1%) positions.

#### Zinc and selenium

Zinc and Se concentrations in soybean seeds were affected differently by the environmental treatments imposed in the present study (Figure 6). Elevated [CO_2_] decreased seed [Zn], but the decrease was only significant in 2015 (−8.9%). In contrast, seed [Zn] was significantly higher under elevated temperature in 2014 (+3.4%) and 2015 (+6.4%). Seed [Zn] decreased towards the top of the canopy in 2014 (top *minus* bottom: −5.6%) and 2015 (top *minus* bottom: −7.7%), and this effect was influenced by temperature, with a more pronounced decrease under the control temperature treatment (top *minus* bottom: 2014, −11.6%; 2015, −12.8%) than under the heat treatment (top *minus* bottom: 2014, 0.3%; 2015, −2.6%) (Table 4). In contrast to Zn, Se is present at substantially lower concentrations and no significant main effects of eCO_2_ or elevated temperature were observed. Variation in the range of seed [Se] was much higher in 2014 and there was a significant effect of canopy position on seed [Se] only in 2014 (top *minus* bottom: −25.0%).

**Figure 5:**
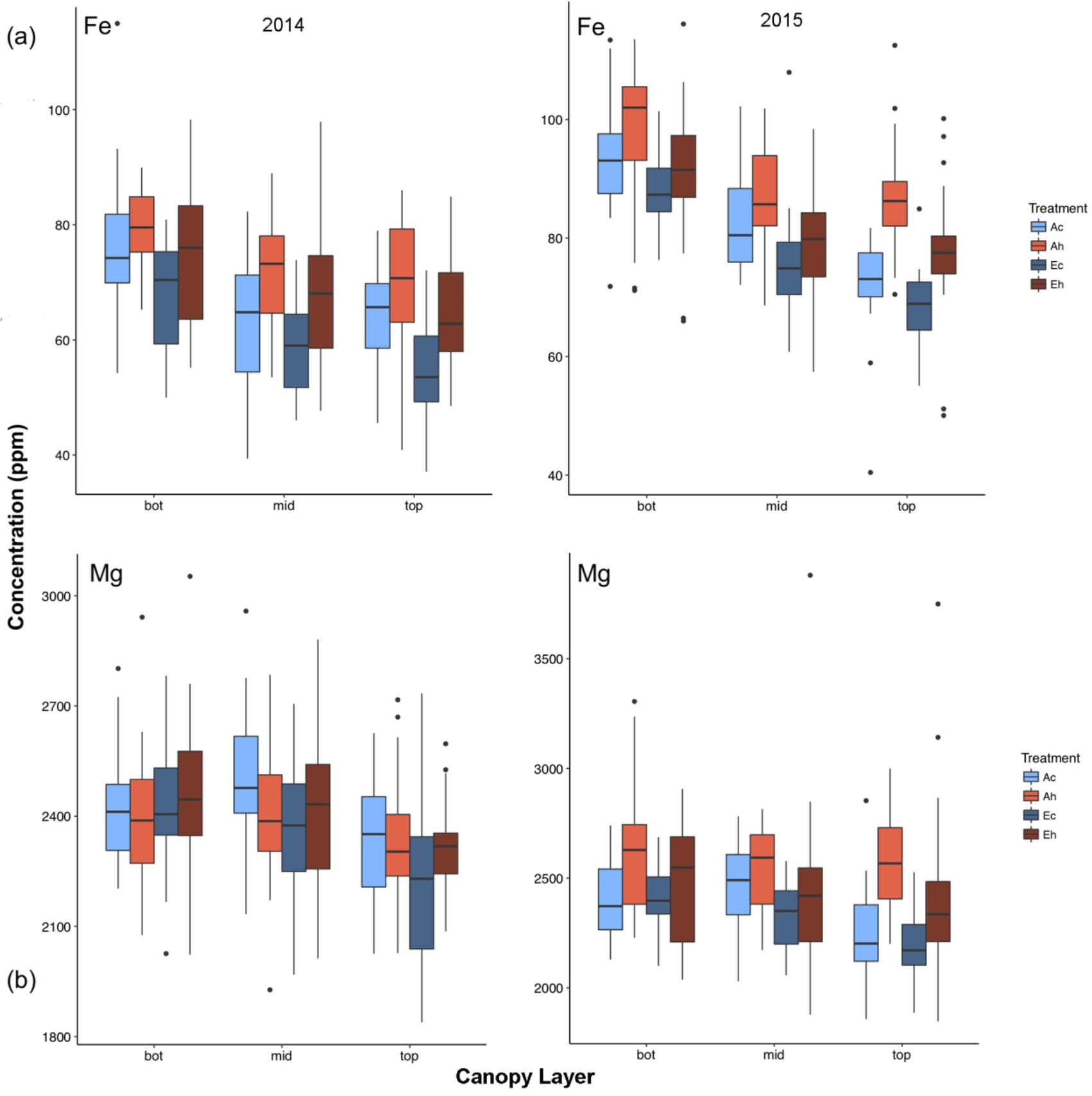
Canopy gradients of (a) seed [Fe] and (b) seed [Mg]. Seed produced at the bottom, middle and top third of the main stem were analyzed from plants grown at eCO_2_ and elevated temperature. Boxplots display the median, and 25 and 75% percentiles, and whiskers extend to 1.5X interquartile range, with outliers (if any) shown. Ac, ambient CO_2_, control temperature; Ah, ambient CO_2_, heated + 3.5 °C; Ec, elevated CO_2_, control temperature; Eh, elevated CO_2_, heated + 3.5 °C.

**Figure 6:**
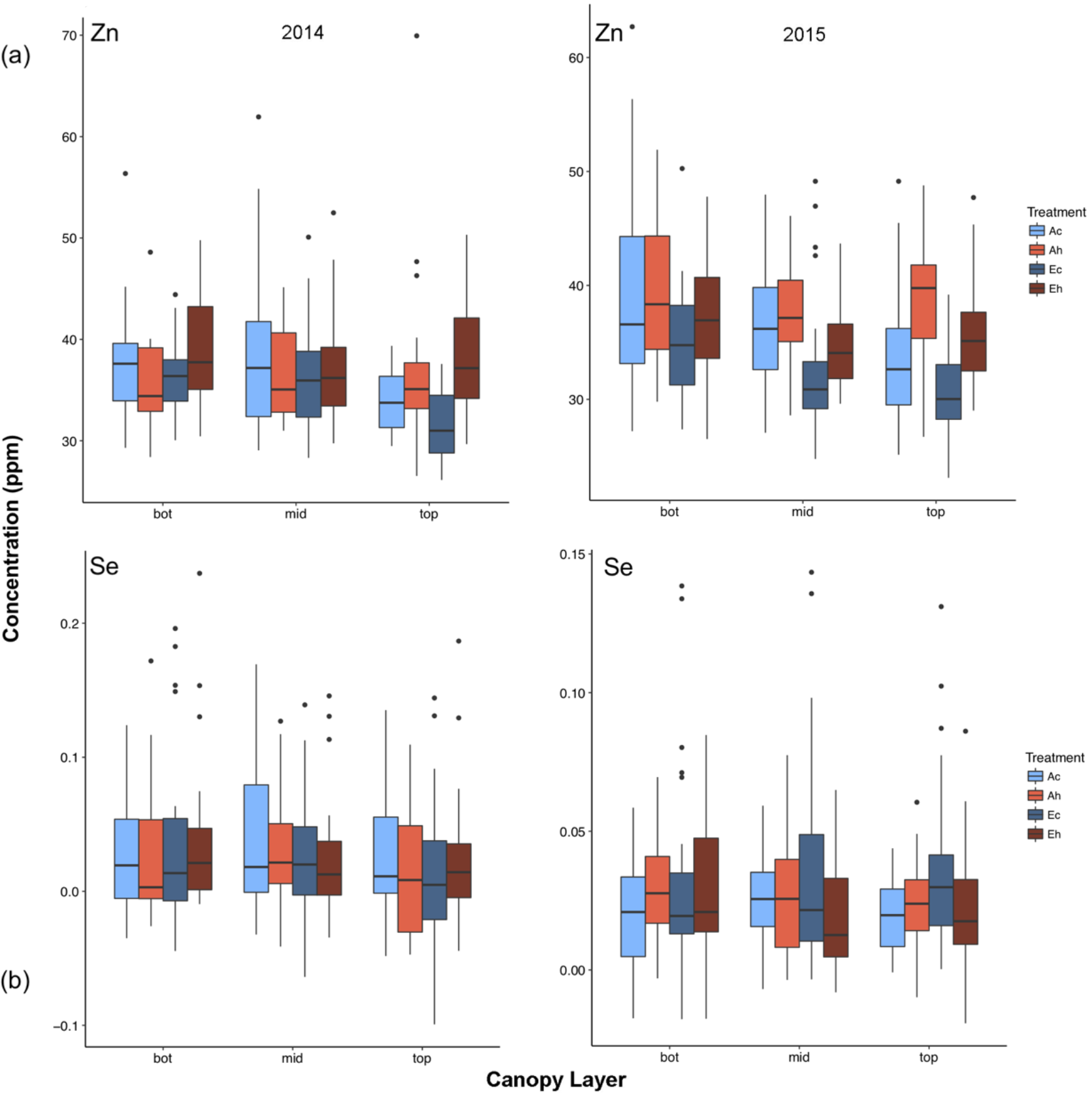
Canopy gradients of (a) seed [Zn] and (b) [Se]. Seed produced at the bottom, middle and top third of the main stem were analyzed from plants grown at eCO_2_ and elevated temperature. Boxplots display the median, and 25 and 75% percentiles, and whiskers extend to 1.5X interquartile range, with outliers (if any) shown. Ac, ambient CO_2_, control temperature; Ah, ambient CO_2_, heated + 3.5 °C; Ec, elevated CO_2_, control temperature; Eh, elevated CO_2_, heated + 3.5 °C.

#### Calcium, manganese and strontium

Calcium, Mn and Sr, which is included in the group because it is a chemical analog of Ca and therefore frequently correlated (Baxter, 2009), are present at very different concentrations in seeds (Ca ≫ Mn > Sr). However, they responded similarly in terms of variation with canopy position, and response to CO_2_ concentration and temperature (Figure 7). For example, the seed concentrations of Ca, Mn and Sr were not affected by eCO_2_, with the exception that seed [Ca] was reduced by 5.5% in just one year (2014). Conversely, elevated temperature increased seed [Ca] and [Sr] in both years but [Mn] only in 2015. With calcium, seed concentrations were significantly higher under the heat treatment in 2014 (+9.1%) and 2015 (+17.9%). Similarly, heat treatment increased seed [Sr] by 11.2% in 2014 and 19.2% in 2015, while for Mn the increase in 2015 was only 5.8%. As noted above, seed [Ca] increased significantly towards the top of the canopy in 2014 (top *minus* bottom position: +29.2%), but in 2015 this effect was mainly observed in the heat treatment (top *minus* bottom: +16.9%) and not for the control temperature treatment (top *minus* bottom: +3.2%). A significant increase of Mn towards the top of the canopy was observed in both seasons (top minus bottom: +26.0% in 2014, +30.9% in 2015). Sr also increased significantly towards the top of the canopy in 2014 (top *minus* bottom position: +74.0%) and to a lesser extent in 2015 (top *minus* bottom position: +38.4%).

#### Boron

At ambient temperature, seed [B] tended to decrease from the bottom to the top of the canopy (top *minus* bottom: 2014, −8.5%; 2015, −11.4%). Elevated CO_2_ significantly decreased [B] by 6.0% in 2015, but this effect was not observed in 2014. In contrast, elevated temperature significantly increased seed [B] in both years (2014, 19.8%; 2015, 17.0%) and there was a significant temperature by canopy interaction. As noted above, seed [B] tended to *decrease* towards the top of the canopy under the control temperature treatment, but *increased* towards the top of the canopy under the high temperature treatment (top *minus* bottom: 2014, +14.0%; 2015, +9.9%). While seed [B] was increased by elevated temperature at all canopy positions, the greatest impact was observed at the top of the canopy (Figure 8).

**Figure 7:**
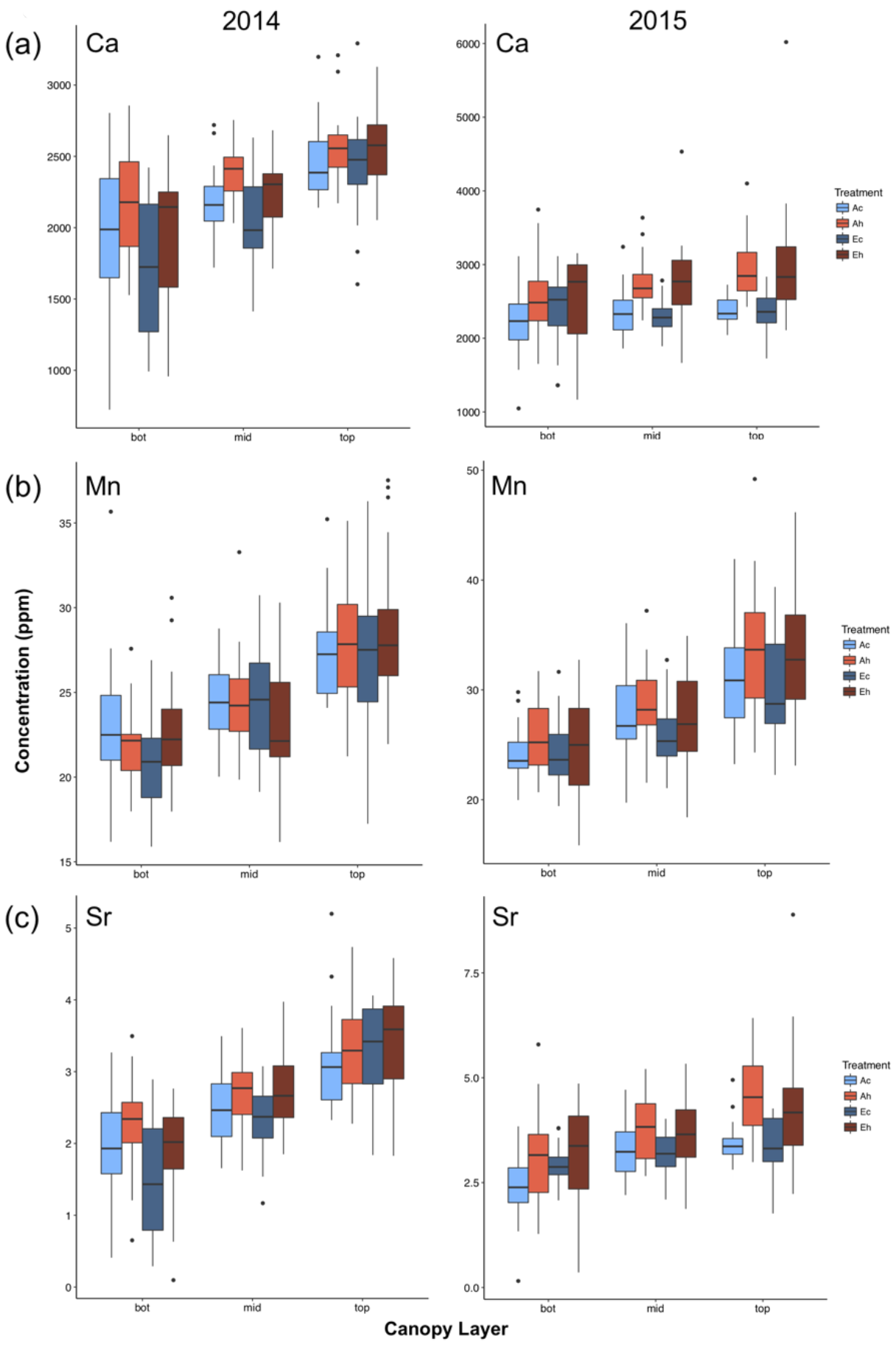
Canopy gradients of (a) seed [Ca], (b) [Mn] and (c) [Sr]. Seed produced at the bottom, middle and top third of the main stem were analyzed from plants grown at eCO_2_ and elevated temperature. Boxplots display the median, and 25 and 75% percentiles, and whiskers extend to 1.5X interquartile range, with outliers (if any) shown. Ac, ambient CO_2_, control temperature; Ah, ambient CO_2_, heated + 3.5 °C; Ec, elevated CO_2_, control temperature; Eh, elevated CO_2_, heated + 3.5 °C.

#### Correlations between seed mineral concentrations and seed yield

In order to determine whether changes in seed yield influenced seed mineral concentrations, we compared seed mineral concentrations with seed yields from the corresponding canopy position and results are summarized in Table 6. With a few exceptions where there was a positive correlation (i.e., one year of study each for P and S), in the majority of cases the correlations were not strong and were not statistically significant (p < 0.10). However, two exceptions emerge with seed [B] and [Fe], where significant negative correlations were observed in both years of the study (Table 6 and Figure 9). These results suggest that B, Fe, and perhaps to a lesser extent Mg and Ca, are in limited or fixed supply to the developing seeds, and that changes in concentrations will reflect, in part at least, changes in number of seeds produced in a given environment.

## Discussion

In the present study, we analyzed the impact of eCO_2_ and elevated temperature, singly and in combination, on soybean seed yield and seed protein, oil and mineral concentrations. The major conclusions are that eCO_2_ increased seed yield and reduced the concentrations of a few minerals such as Fe, whereas elevated temperature reduced yield and increased the concentration of a range of minerals. In general, elevated temperature counteracted the increase in yield and the reduction of mineral concentrations (where observed), suggesting that the previously reported increase in seed yield and threat to human nutrition associated with eCO_2_ (Loladze, 2014, McGrath and Lobell, 2013, Myers et al., 2014) may not be realized when combined with elevated temperatures (Ruiz-Vera et al., 2013, Köhler et al., 2016). These conclusions are discussed in more detail below.

**Table 6:** Overview of correlation analysis between seed mineral concentration and total seed yield at the corresponding canopy position. Where statistically significant (p < 0.10) the p-values are shown parenthetically.

| Element | 2014 | 2015 |
| --- | --- | --- |
|  | r value |  |
| B | *-0.52 (p = 0.00017) | *-0.36 (p = 0.012) |
| Na | ns | ns |
| Mg | ns | *-0.25 (p = 0.089) |
| Al | ns | ns |
| P | *0.51 (p = 0.00021) | ns |
| S | ns | *0.33 (p = 0.024) |
| K | ns | *-0.27 (p = 0.059) |
| Ca | *-0.40 (p = 0.0047) | ns |
| Fe | *-0.37 (p = 0.0009) | *-0.60 (p = $7.3 \times 10^{-6}$) |
| Mn | ns | ns |
| Co | ns | ns |
| Ni | *-0.26 (p = 0.075) | ns |
| Cu | ns | ns |
| Zn | ns | ns |
| As | ns | *-0.36 (p = 0.011) |
| Se | ns | ns |
| Rb | ns | ns |
| Sr | *-0.30 (p = 0.04) | ns |
| Mo | ns | ns |
| Cd | *-0.34 (p = 0.018) | ns |

**Figure 9:**
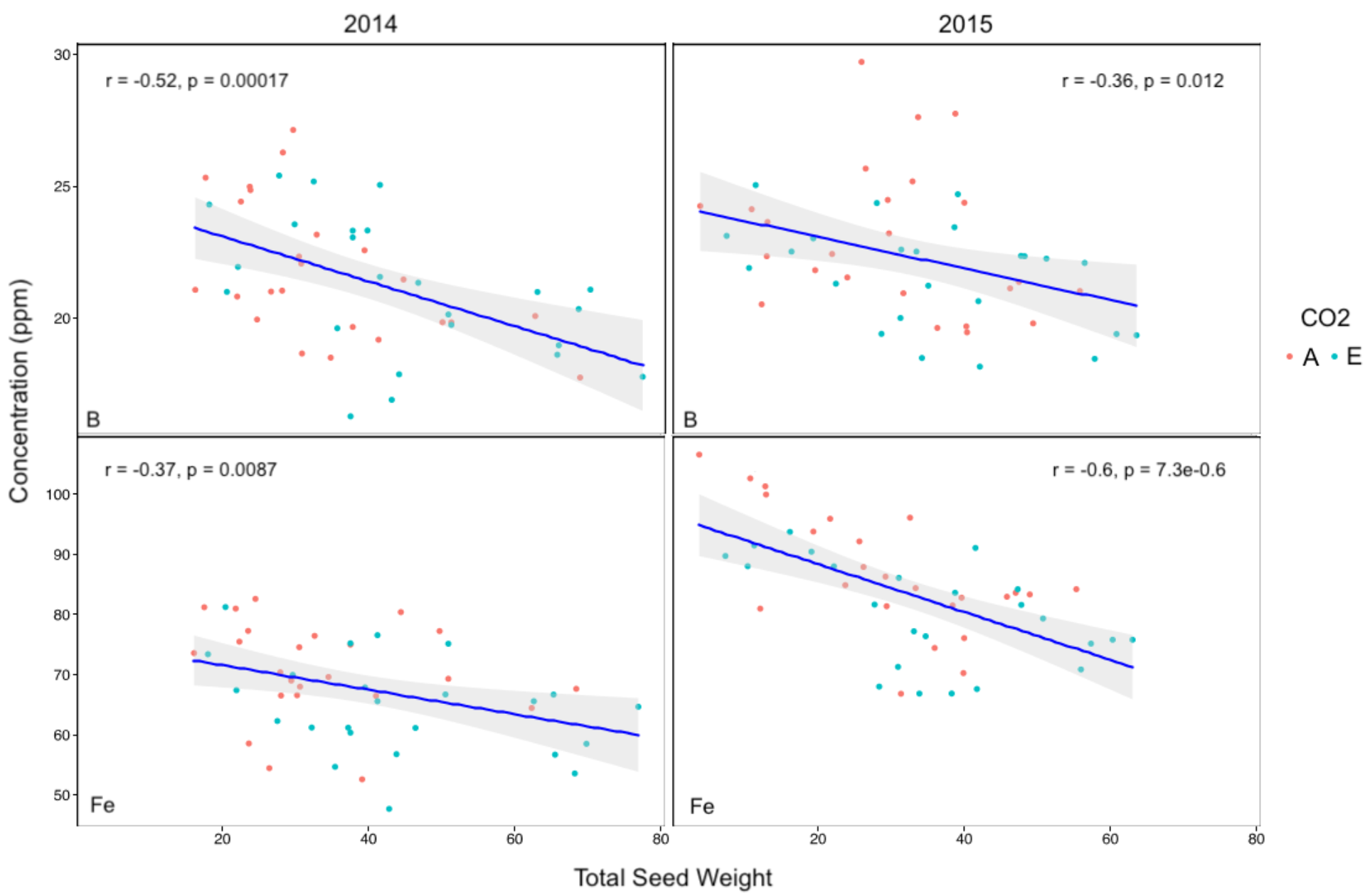
Correlation plots between seed mineral concentration and seed yield at the corresponding canopy position for (a) seed [B] and (b) seed [Fe]. Data points are color coded for samples taken from plants grown at ambient (red) and eCO_2_ (blue).

### Elevated temperature reduces seed yield differentially through the canopy

As would be expected (Sionit et al., 1987, Gibson and Mullen, 1996, Huxley et al., 1976, Ruiz-Vera et al., 2013), the elevated temperature treatment reduced seed yield on both a land area basis (Figure 2) and per plant basis (Figure 3a). However, looking at seed yield as a function of position within the canopy suggested that elevated temperature preferentially reduced yield in the bottom third of the canopy (Figure 3b). This is somewhat counter-intuitive as it would be reasonable to assume that elevated temperature generated by infrared heaters (that do not increase air temperature) would primarily have impact in the upper portion of the canopy. However, the observed result would be consistent with the notion that upper canopy leaves play the primary role in provision of assimilates (reduced-C in the form of sucrose and reduced-N in the form of amide amino acids) to support seed development throughout the canopy; when leaf photosynthesis (and hence assimilate supply) is limited by elevated temperature (Ruiz-Vera et al., 2013), seed development lower in the canopy would be most attenuated. The unexpected impact of the heat treatment on yield of seed produced at the bottom canopy position was observed at both ambient and eCO_2_.

### Elevated temperature, but not eCO_2_, affects seed protein and oil concentrations

Consistent with earlier reports (Escalante and Wilcox, 1993b, Escalante and Wilcox, 1993a, Collins and Cartter, 1956, Huber et al., 2016), there was a significant canopy position effect on seed storage product concentration, with seeds at the top of the canopy having higher protein and seeds at the bottom having higher oil concentrations (Figure 4). In general, eCO_2_ did not have a significant effect on seed protein or oil concentrations, as observed in other studies with soybeans (Jin et al., 2017, Myers et al., 2014, Bishop et al., 2015). In contrast, elevated temperature tended to reduce seed protein concentration and increase seed oil concentration, which may be related to the previously reported reduction of leaf photosynthesis at high temperature (Ruiz-Vera et al., 2013) that could impact seed composition throughout the canopy.

### Impact of eCO_2_ and elevated temperature on seed minerals

Consistent with previous reports, elevated CO_2_ reduced the concentration of Fe and Zn significantly (with the exception of Zn in 2014). No other elements were significantly impacted by eCO_2_, although the general trend was to reduce mineral concentrations, consistent with earlier reports (Loladze, 2014, McGrath and Lobell, 2013, Myers et al., 2014, Weigel, 2014). In contrast, elevated temperature significantly affected the concentrations of Fe, Zn and a number of minerals not affected by eCO_2_. Thus, elevated temperature generally had the opposite effect of eCO_2_. These changes suggest that there is a linkage between the increased seed yield and decreased seed mineral concentrations in eCO2, and that elevated temperature reverses both effects. Collectively, these results suggest that the previously reported effects of increasing atmospheric [CO_2_] on seed mineral concentration may not be realized in future climates where eCO_2_ occurs with other environmental changes such as elevated temperature. Interestingly, the observed effects of elevated temperature and eCO_2_ on seed minerals does not correlate with the mode of mineral delivery and uptake into the plant; i.e., both treatments affected minerals that move by mass flow and are therefore affected by transpiration (e.g., Ca, Mg, and S), as well as those with diffusion to the root as the main mechanism (e.g., K, P, Mn, Zn, Cu and Fe) (Oliveira et al., 2010, Oliver and Barber, 1966).

To accumulate in seeds, minerals need to be available in the soil water; presented to the root surface by mass flow or diffusion; taken up into the root; distributed within the plant; and finally partitioned to the developing seeds. Limitations at any of these steps could result in changes in seed mineral concentrations when the number of seeds produced is altered and seed size does not change (as was observed in this study). For example, a mineral present in limiting or fixed amounts may be reduced in concentration in seeds when the number of seeds produced is increased (e.g., at eCO_2_) and conversely increased in concentration when the number of seeds produced is decreased (e.g., at high temperature). In such a case, seed mineral concentration would be negatively correlated with seed yield, reflecting enrichment when yield was reduced and dilution when growth was increased. Indeed, this was observed for seed [B] and [Fe] in both years of the study, and in one year of study for seed [Ca] and [Mg]. Exactly which step in the process of mineral accumulation in the developing seed is limiting or highly regulated may vary with the mineral, and will be an important focus for future research. Fundamentally, this study suggests that the effects of climate change will be variable on nutrient and caloric levels, and our current knowledge is insufficient to predict the results.

### The unusual response of B to eCO_2_ and elevated temperature

Boron is an essential micronutrient and while the functions of this micronutrient are not fully understood it is recognized that B may impact various aspects of plant metabolism and transport (Pilbeam and Kirkby, 1983) and cell wall composition (O’Neill et al., 2004). In terms of transport, B is present at neutral pH primarily as boric acid, which can be transported along with the borate anion (Woods, 1996). However, B can be more readily transported as a B-polyol complex (Brown and Shelp, 1997). It is thought that species that produce polyols (e.g., sugar alcohols such as sorbitol and mannitol) may display greater B mobility within the plant, in particular in the phloem, and may explain the large variation in B mobility among species that has been noted. One intriguing possibility is that at elevated temperature, soybean leaves produce more sugar alcohols compared to ambient temperature, which could result in the increased seed [B] observed. This speculative mechanism would also be consistent with increased seed [B] at the top rather than bottom of the canopy (Figure 8). This is clearly an area for further study.

## Concluding Remarks

The reduction in a broad range of nutritionally important minerals in soybean seeds as a result of growth at eCO_2_ (Loladze, 2014, McGrath and Lobell, 2013, Myers et al., 2014) was generally confirmed in the present study, and conversely, the increase in seed mineral concentrations by growth at elevated temperature (at ambient CO_2_) was shown. In general, growth at eCO_2_ plus elevated temperature restored seed yield and mineral concentrations to nearly those observed at current ambient conditions, calling into question the predictive value of studies looking only at eCO_2_ to identify threats to human nutrition. The general inverse relationship between mineral concentration and seed yield per plant as affected by eCO_2_ and elevated temperature, suggests that dilution by increased growth (at eCO_2_) or enrichment by reduced growth (at elevated temperature) may be general factors impacting seed mineral concentrations. It remains to be determined how broadly applicable our results are outside of the single cultivar of soybeans used in this experiment. Other cultivars and species may respond differently to either or both stresses, with unknown effects on the nutritional components. We are also unable to predict whether breeding programs conducted in these predicted conditions will be able to ameliorate the loss of nutritional content while maintaining the yield gains.

**Figure 8:**
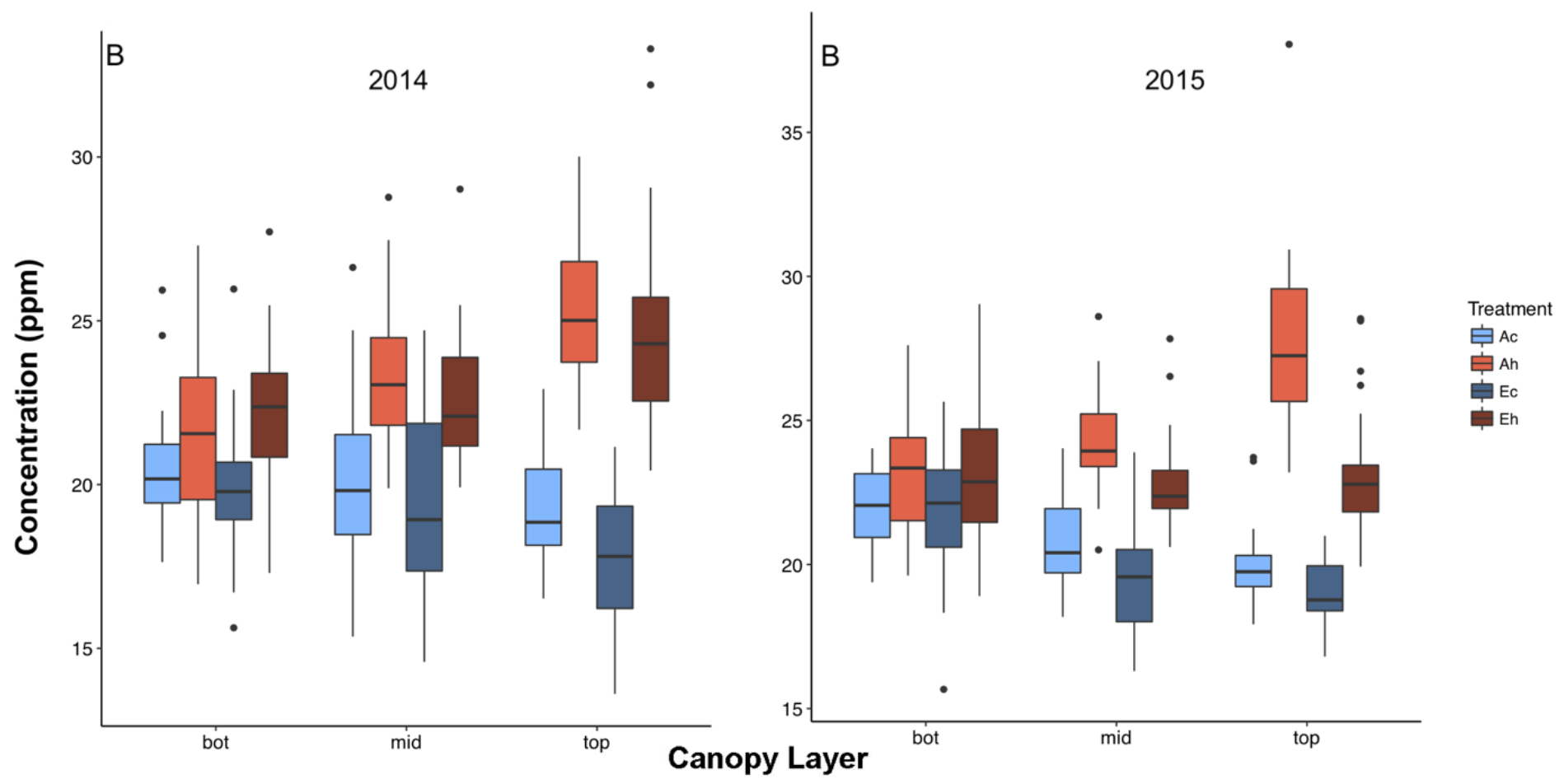
Canopy gradients of seed [B]. Seed produced at the bottom, middle and top third of the main stem were analyzed from plants grown at eCO_2_ and elevated temperature. Boxplots display the median, and 25 and 75% percentiles, and whiskers extend to 1.5X interquartile range, with outliers (if any) shown. Ac, ambient CO_2_, control temperature; Ah, ambient CO_2_, heated + 3.5 °C; Ec, elevated CO_2_, control temperature; Eh, elevated CO_2_, heated + 3.5 °C.

Another noteworthy point is that seed composition of protein, oil and some minerals varies with node position at which the seeds developed, and confirms our earlier study (Huber et al., 2016). Moreover, environmental factors that affect composition tend to impact seed produced at all positions within the canopy, even though the effects on yield are not necessarily uniform across the canopy. The largest exception to this generalization was seed [B], which was strikingly increased in seeds at the top of the canopy when plants were grown at elevated temperature. As discussed above, the potential for altered metabolism at elevated temperature resulting in enhanced B mobility within the plant is an attractive hypothesis for further testing.

## Experimental Procedures

### Plant Material and Growth Conditions

Soybean [*Glycine max* (L.) Merr. cv. ‘Thorne’] was grown in a complete block design (*n* = 4) in the Soybean Temperature by Free Air CO_2_ Enrichment (Soy-T-FACE) experiment at the SoyFACE field site near Urbana-Champaign, Illinois, USA (40°2’30.49” N, 88°13’58.80” W, 230 m above sea level) during 2014 and 2015 growing seasons. The experiment consisted of four blocks, each containing one ambient and one eCO_2_ plot and within each plot there was an unheated and a heated sub-plot. Seeds were planted by hand at 5-cm intervals in 38-cm rows. Eight 11-m long rows were planted in each of the eight plots (four rows of the wild-type ‘Thorne ’ used in this study, and the other four of a transgenic soybean line that was used for another experiment). The ambient [CO_2_] plot was at approx. 400 μmol mol^−1^ and the elevated one at approx. 600 μmol mol^−1^, using free air CO_2_ enrichment (FACE) fumigation technology (Miglietta *et al,* 2001). The heated sub-plots were each equipped with an infrared heater array, as described in detail previously (Ruiz-Vera *et al,* 2013) installed at 1.0 m to 1.2 m above the canopy on a telescoping mast system (Ruiz-Vera *et* al., 2015). Using a PID feedback control system we warmed the crop canopy to a target elevation of +3.5 °C above that of the canopy temperature in the unheated sub-plot. The target temperature increase was based on the low-response model predictions for surface temperature in the Midwest in 2050 (Rowlands *et al,* 2012). During the day (6 am to 6 pm) and with rainy days excluded, mean temperature differences between the sub-plots were between 0.5 and 1.0 °C lower than the target set point (Table 1), resulting in an average temperature increase of +2.7 °C. During the night the average temperature difference was +3.4 °C. The heated sub-plot diameter was 3.5 m resulting in an effective heated sub-plot area of 9.6 m^2^. Canopy temperature in each sub-plot was measured by infra-red radiometers (SI-111; Apogee Instruments) connected to data-loggers CR1000 Micrologger, Campbell Scientific, Logan, UT, USA). Canopy temperature measurements were collected every 5 seconds to control the heater output, averaged every 10 min, and these values were stored. For more information, see Köhler et al. (2016). In the figures, the treatments are referred to as ‘*Ac’* (**A**mbient [CO_2_], **c**ontrol temperature), ‘*Ah*’ (**A**mbient [CO_2_], **h**eat), ‘*EC*’ (**E**levated [CO_2_], **c**ontrol temperature) and ‘*Eh*’ (**E**levated [CO_2_] + **h**eat).

### Harvest and Seed Yield Determination

After full maturity and dry-down was complete (R8), 10 plants were harvested by hand (stems were cut with scissors at about 2.5 cm above the ground), wrapped in burlap bags and air dried at 27 °C in a drying room. The individual plants were cut into three equal parts (bottom, middle and top sections) and the pods from the three sections threshed separately and the seeds weighed. The three sections are hereafter referred to as bottom middle, and top canopy positions.

### Protein and Oil Analysis

Protein and oil were measured with an Infratech 1241 Grain Analyzer (FOSS Analytical AB, Höganäs, Sweden), which is a true Near Infrared Transmission instrument that generates a spectrum from 850 to 1050 nm via the monochrome light source and mobile grating system. A 50-ml seed sample was used that allowed for 10 subsample readings reported on a 13 % moisture basis.

### Mineral Analysis

Six seeds from each sample were selected at random from the seed yield samples for mineral analysis as described in (Ziegler et al., 2013). In brief, single seeds from each main stem canopy position were weighed using a custom-built seed weighing robot and then digested in concentrated nitric acid before loading onto an Elan ICP-MS. Internal standards were used to control for differences in dilution and sample injection. Custom scripts were used to correct for internal standards and correct for sample weight.

### Statistical Analysis

Data were analyzed using a randomly blocked mixed-model analysis of variance (PROC MIXED, SAS 9.4) taking into account the split-plot design of the experiment. The analyses were done separately for individual years. The model for the seed yield, protein and oil data from harvest of individual plants and for the elemental concentrations included the fixed factors [CO_2_] (ambient, elevated), temperature (control, heated), and position in the canopy (bottom, middle or top). The model for the bulk seed yield included only the fixed factors [CO_2_] (ambient, elevated) and temperature (control, heated). Block was included as a random factor in all five models. For the analysis of the elemental concentrations, the results from the individually analyzed six seeds per sample were averaged for each canopy layer within each subplot. Significant differences between least square means for *a priori* determined comparisons were analyzed using *post hoc* tests (LSMEANS). Probability for statistical significance was set at p < 0.1 *a priori* to reduce the possibility of type II errors. All data and analysis scripts used in the analysis are available on www.ionomicshub.org.

## Acknowledgements

This work was supported by the USDA-Agricultural Research Service. This material is based upon work that is supported by the National Institute of Food and Agriculture, U.S. Department of Agriculture, under award number 2014-67013-21783. The authors thank Greg Ziegler for skilled technical assistance in the mineral analysis and Jennifer Barrett and Amber Wolf for assistance with figure creation.

---

**Figure S1.**
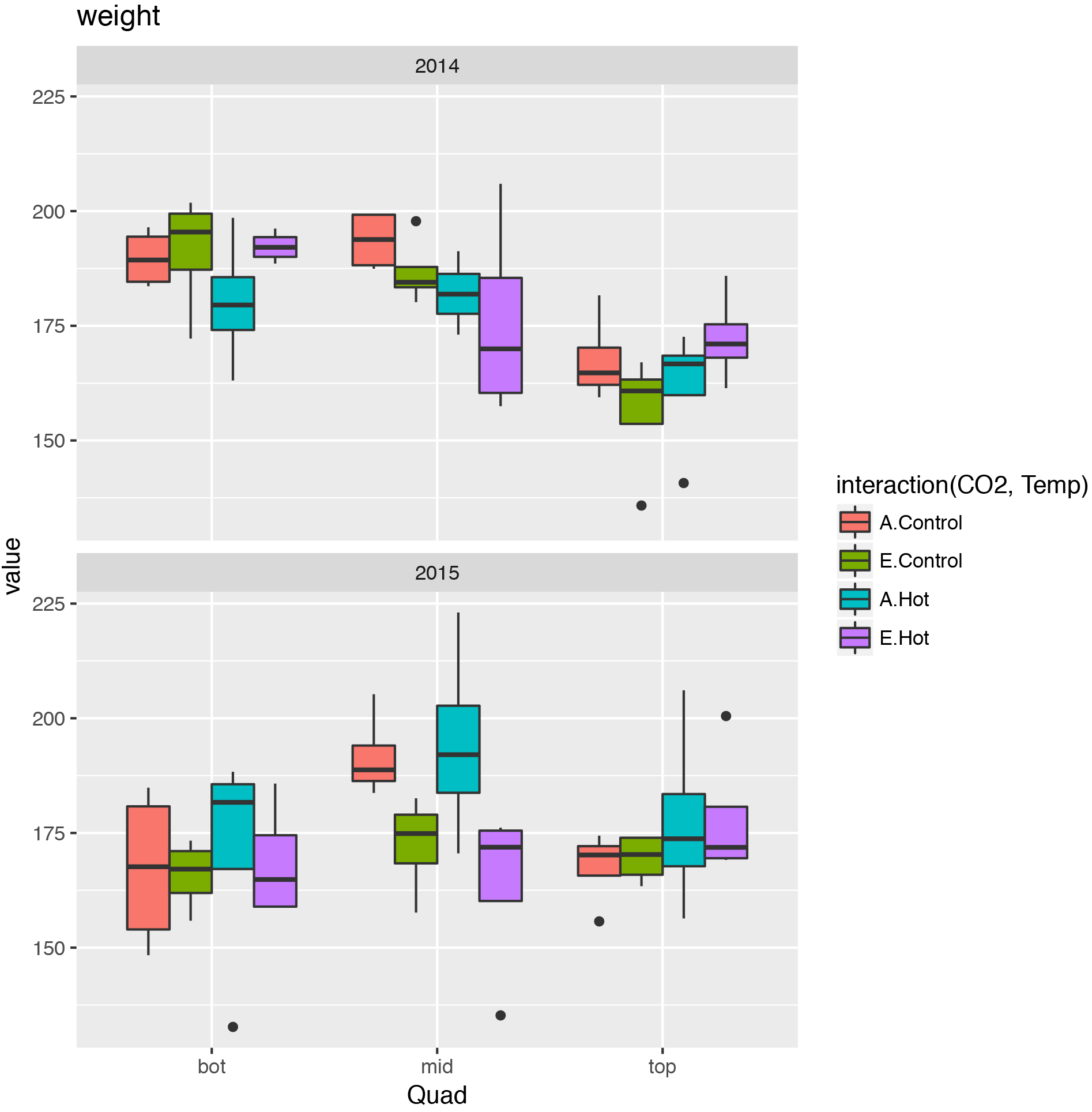
Variation in single seed weight as a function of canopy position (bottom, middle or top third of the main stem) where the seeds were produced. Ac, ambient CO_2_, control temperature; Ah, ambient CO_2_, heated + 3.5 °C; Ec, elevated CO_2_, control temperature; Eh, elevated CO_2_, heated + 3.5 °C. Error bars represent ±1 standard error of the estimate (LSM).

**Figure S2.**
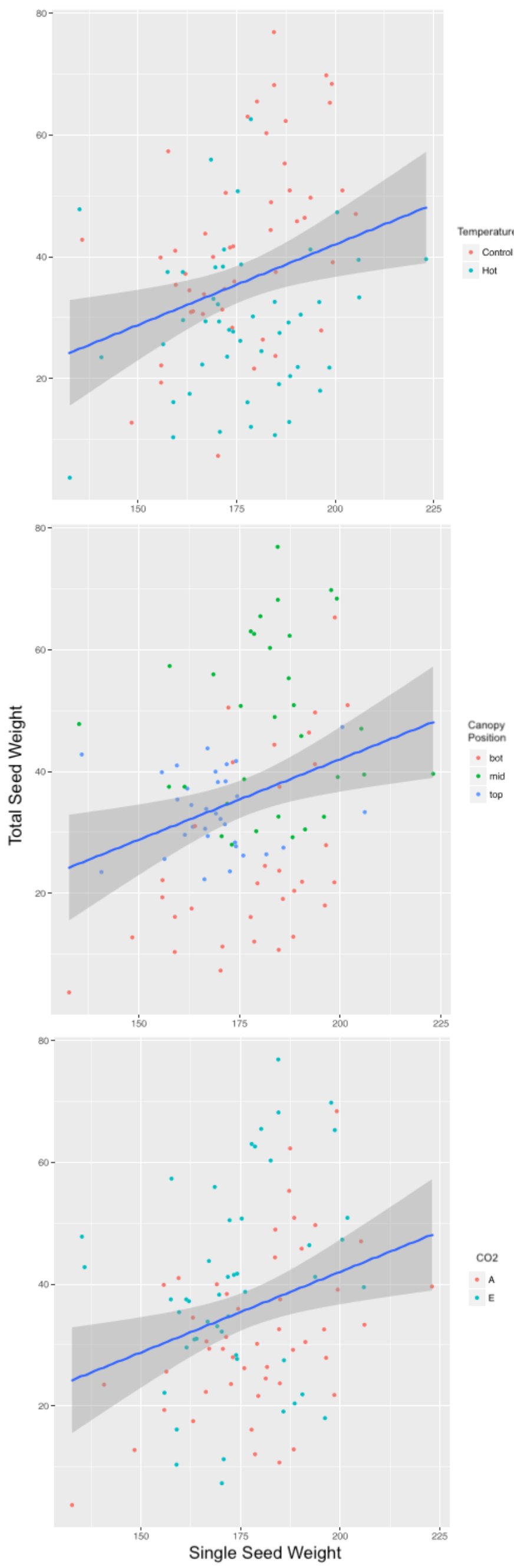
Correlation plots between single seed weight and seed yield for the corresponding canopy position. Results from 2014 and 2015 were combined and data points are color coded according to (a) ambient vs elevated temperature; (b) canopy position; and (c) ambient vs elevated CO_2_. The correlations were not statistically significant (p = 0.28) and simply document that changes in yield were primarily driven by variation in number of seeds produced.

